# A global study of the geographic range size of epiphytes

**DOI:** 10.1101/2023.11.20.567933

**Authors:** Vida J. Svahnström, Eimear Nic Lughadha, Félix Forest, Tarciso C.C. Leão

## Abstract

Epiphytes have long been considered to have larger geographic range sizes than terrestrial plants, yet evidence for this claim comes from studies at restricted geographic and taxonomic scales and is contrary to that of some recent studies. We examined if epiphytes have larger or smaller range sizes than terrestrial plants and tested if epiphytism is a likely driver of differences in range size globally across angiosperms. We integrated global datasets on angiosperm taxonomy, distribution, and lifeform to calculate three range size metrics. We tested if there were significant differences in mean range size between epiphytes and terrestrial plants across angiosperms and within epiphyte-rich families using ordinary and phylogenetic regression models. On average, epiphytes have larger range sizes than closely related terrestrial species, supporting the hypothesis that epiphytism favours dispersal into larger areas. However, species in families where epiphytism is prevalent tend to have small range sizes regardless of their lifeform. A high proportion of epiphytes and their close relatives are rare or have vulnerably small range sizes, yet epiphytism per se does not cause rarity. Evolutionary histories and shared traits of epiphyte-rich lineages likely underlie the observed rarity and small ranges.

## Main

The area over which species are distributed, their geographic range size, is an important but poorly-understood factor underpinning global plant diversity and vulnerability to extinction^1,2^. The probability of threats affecting a species’ entire population, thus increasing their risk of extinction, increases as range size decreases^3^. Understanding drivers and correlates of small range size will enhance extinction risk prediction and address urgent needs for species and area prioritisation for conservation.

Epiphytes, plants that germinate and root non-parasitically on other plants^4^, comprise 8% of global angiosperm species^5^, ≤20-39% of total flora in tropical plant diversity centres^6^, and >50% of species in local inventories in certain tropical regions^7,8^. Epiphytes provide critical arboreal habitats^9,10^ and contribute to water and nutrient cycles in tropical forests^11^. Epiphytism occurs in nearly 60 angiosperm families, but epiphyte diversity is concentrated in just a few species-rich lineages: orchids alone comprise 75% of angiosperm epiphyte species^5^.

Despite long-held views that epiphytes have larger ranges than terrestrial species^12,13^, published work suggests a more complex scenario. Andean studies report epiphytes in Bromeliaceae, Piperaceae, and Araceae having larger ranges than confamilial terrestrial species while epiphytic orchids have smaller mean ranges than terrestrial orchids^14,15^. Similar patterns occur at finer taxonomic scales. In the orchid genus *Galeandra,* epiphytic species tended towards smaller range size than terrestrial counterparts^16^. In contrast, epiphytic *Anthurium* species had mean range size eight times larger than terrestrial species in the genus^17^. Analyses of angiosperms in Brazil’s Atlantic Forest showed epiphytes having among the highest percentages of small-ranged species among lifeforms^1,18^.

Adaptations for tree-to-tree dispersal predispose epiphytes to high dispersal capabilities^15^, contributing to large range sizes^19^. Tree height favours dispersal of wind-, bird- or bat-dispersed seeds of canopy-dwelling epiphytes^17,20^. Conversely, plant clades which have undergone rapid diversification tend to have small-ranged species, especially when speciation mechanisms isolate small populations^21,22^. Rapid evolutionary diversification in speciose epiphytic clades could therefore lead to small ranges^23^.

Existing studies indicate that relationships between epiphytism and range size vary between lineages and regions, such that inferring global patterns of range size from local or regional patterns is problematic. Local or regional range size studies occur within particular biogeographical contexts where regionally-specific extrinsic factors could differentially impact terrestrial and epiphytic plants, for example historical climatic changes^24^. Previous studies also failed to capture significant portions of the taxonomic diversity of epiphytes. Moreover, most earlier analyses did not account for range size heritability^25–27^, potentially masking relationships between epiphytism and range size by failing to control for the non-independence of species range sizes caused by shared evolutionary history^28^.

Here, we integrate multiple data sources for near-comprehensive taxonomic and geographic coverage to investigate range size of epiphytes at globally- and taxonomically-inclusive scales for the first time. We test if epiphytes differ in geographic range size from terrestrial plants, across angiosperms and in epiphyte-dominated lineages. Applying phylogenetic comparative methods, we test if epiphytism is a likely driver of observed range size differences. To inform conservation, we also use non-phylogenetic statistics to estimate epiphyte rarity and range-based vulnerability within the broader context of angiosperms.

## Results

We used three metrics to investigate if epiphytes differ in range size from terrestrial plants. First, we calculated the number of botanical countries in which a species is native for all angiosperm species; ‘botanical country’ is the standard unit used to record plant distributions in the World Checklist of Vascular Plants (WCVP)^29,30^. Two-thirds (65%) of epiphyte angiosperms are native to a single botanical country. The epiphyte with the largest range size, the pantropical orchid *Polystachya concreta*, is native to 81 botanical countries. In contrast, 56% of terrestrial angiosperms are native to a single botanical country, and the most wide-ranged terrestrial angiosperm, the cosmopolitan grass *Phragmites australis,* is native to 213 botanical countries. Angiosperm epiphytes were absent from 72 botanical countries at high latitudes or consisting mostly of drylands.

As most variation in range size not captured by number of botanical countries is among small-ranged species, particularly those occurring in 4 or fewer botanical countries, we calculated two additional metrics of range size for angiosperms native to ≤4 botanical countries: sum of unique herbarium specimen records for a species (hereafter ‘specimen count’) and extent of occurrence (EOO). After data cleaning and name-matching with the phylogenetic tree^31^ our dataset contained specimen counts for 72.7% of angiosperms and EOOs for 43.4% of angiosperms. Specimen counts for epiphytes native to ≤4 botanical countries ranged from one to 1,224 with a median of five records after cleaning (Fig. 1). For terrestrial plants, specimens counts ranged from one to 4,048 with a median of 14 records. The EOO of epiphytes native to ≤4 botanical countries ranged from 0.01 km^2^ to 7,431,335 km^2^ (median 29,003 km^2^). For terrestrial plants, EOO ranged from 0.002 km^2^ to 10,696,304 km^2^ (median 45,472 km^2^).

**Figure 1:**
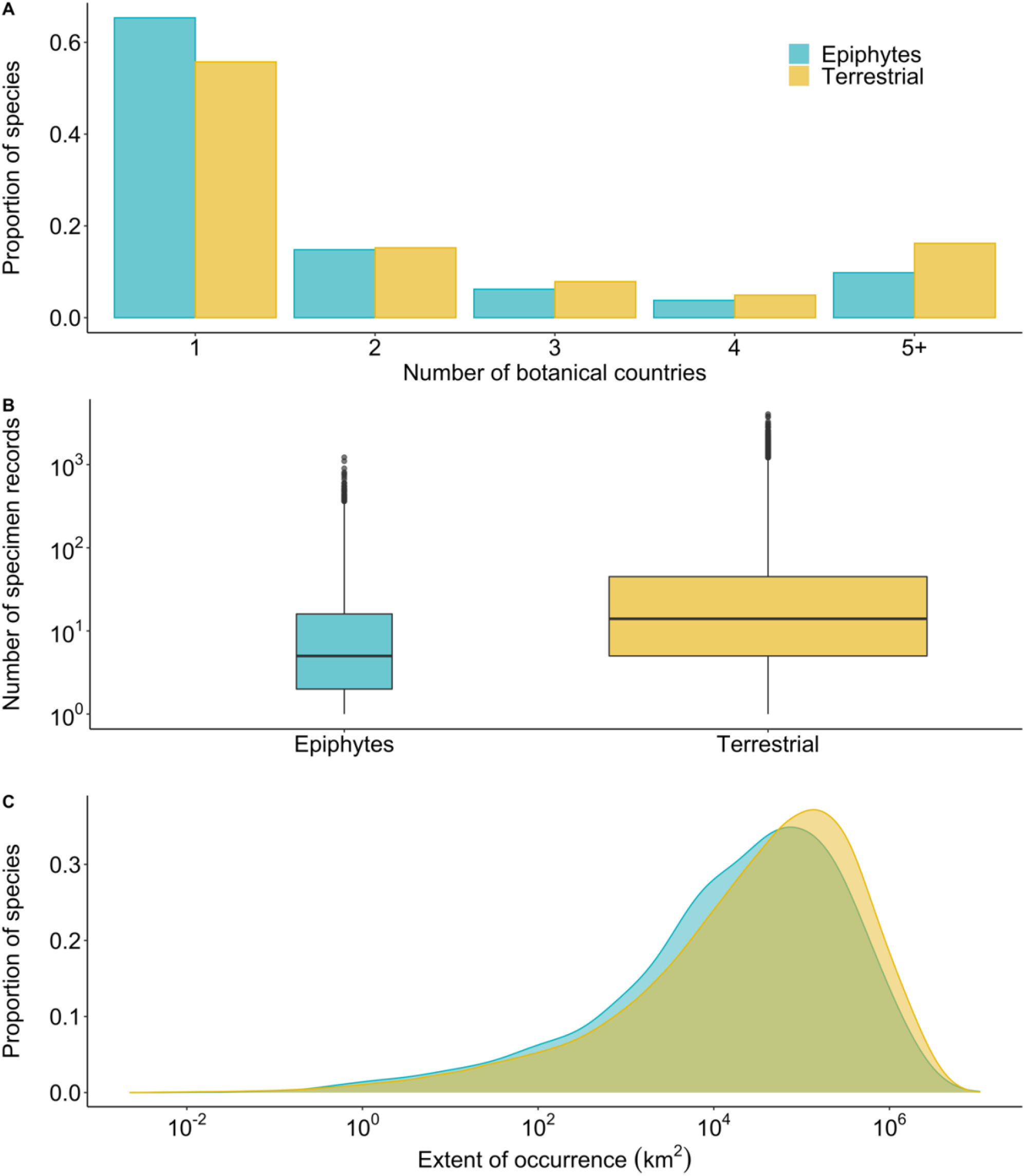
Epiphytes tend to have smaller average range size and larger proportions of species with small ranges, but similar overall distributions of range size to terrestrial plants across three different metrics: A) number of botanical countries, B) number of specimens, and C) extent of occurrence (EOO). The barplot in A shows the proportion of epiphytes and terrestrial plants found in 1-5+ botanical countries. Boxplots in B show the minimum, first quartile, median, third quartile, maximum, and outliers for the number of specimens transformed at the log_10_ scale, with boxes proportional to the number of species in each group. C shows the frequency distributions of the extent of occurrence for epiphytes and terrestrial plants transformed at the log_10_ scale.

The number of botanical countries (1-4) was correlated with specimen count and with EOO at r=0.42 and r=0.49, respectively, supporting the expectation that the number of botanical countries captures a reasonable portion of the variation in range size (even at the low end of range distribution), while EOO and specimen count were correlated at r=0.60 (variables log-transformed, Fig. S1).

### Regression analyses

Across angiosperms, our regression models showed that epiphytes had significantly smaller average range sizes across all three metrics when not controlling for the effect of shared ancestry (range size ∼ lifeform (terrestrial or epiphytic); Fig. 2). Epiphytes are, on average, found in 36% (95% CI: 34-38%) fewer botanical countries, have 58% (95% CI: 57%-59%) fewer specimens and have 33% (95% CI: 30%-36%) smaller EOOs than terrestrial species. The similarity in effect sizes supports using each metric as a credible measure of range size. When restricting analyses to tropical species, effect sizes varied by fewer than 10 percentage points for all three metrics, demonstrating that global patterns across angiosperms are not confounded by the primarily tropical distribution of epiphytes (Fig S2). Excluding angiosperms which occur on oceanic islands had a similarly small effect of less than 6 percentage points across metrics (Fig. S3).

**Figure 2:**
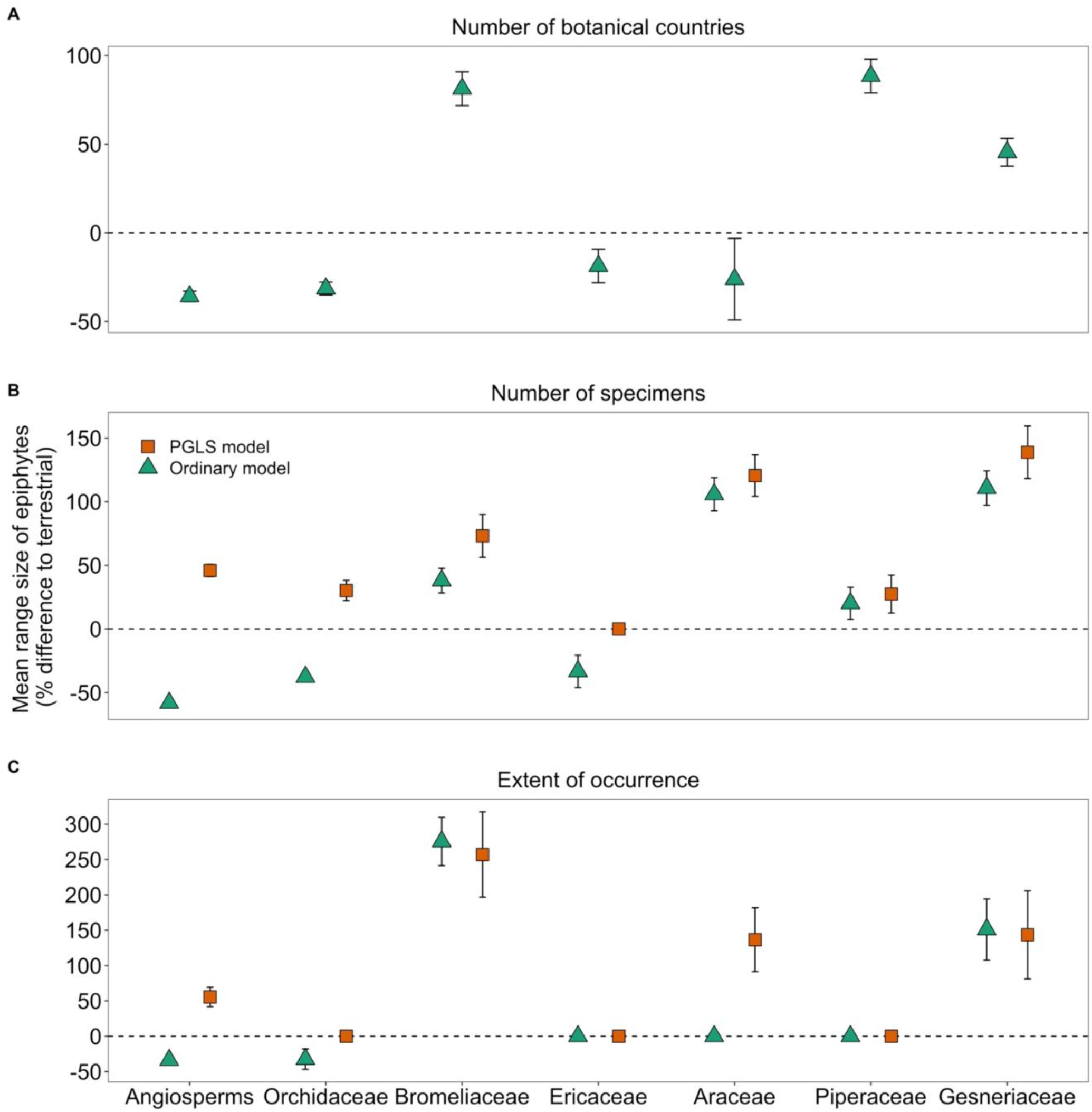
Across angiosperms, epiphytes have smaller mean range sizes than terrestrial plants in ordinary (non-phylogenetic) regression models and larger mean range sizes than terrestrial plants in phylogenetic generalised least squares (PGLS) models across all metrics. However, within-family patterns of range size variation differ between the six most epiphyte-rich families. Mean range size is given as percentage (%) difference to terrestrial species with 95% confidence intervals for the three metrics: A) number of botanical countries (PGLS not performed); B) specimen count; C) extent of occurrence. The horizontal dotted line represents the baseline of no significant difference between epiphytes and terrestrial plants.

Twenty-four angiosperm families include ≥10 epiphytic species (Table S4). Analysis of how range size of lifeforms differs within these families showed considerable variation from the angiosperm-wide trend and between families (Tables S1-S3). Epiphytes showed larger mean specimen count and EOO than confamilial terrestrial species in more cases than the converse. In most families (63%), epiphytes differed from terrestrial species in mean specimen count, but far fewer (25%) differed in their EOOs.

Although 56 angiosperm families contain epiphytic taxa, 92% of epiphyte diversity is concentrated in just six families: Orchidaceae (n=21,034 epiphyte species), Bromeliaceae (n=1,937), Ericaceae (n=876), Araceae (n=766), Piperaceae (n=699), and Gesneriaceae (n=644). Comparison of range size between lifeforms within these families showed that the only family in which epiphytes had fewer specimens and smaller EOOs than terrestrial species was the species-rich, mostly epiphytic Orchidaceae (Fig. 2 (ordinary model)), which contains 75% of angiosperm epiphyte species. In Bromeliaceae and Gesneriaceae, epiphytes had both more specimens and larger EOOs than terrestrial confamilial species. However, these differences are largely driven by the small range sizes of their terrestrial family members, particularly the extremely small ranges of terrestrial bromeliads (Fig. 3). In the remaining epiphyte-rich families, mean specimen count was either larger (Araceae and Piperaceae) or smaller (Ericaceae) for epiphytes, while mean EOO did not differ between lifeforms (Fig. 2). Sensitivity analyses restricted to tropical species yielded similar results, with appreciable differences in effect size only in Orchidaceae and Ericaceae (Fig S2). Epiphytic orchids showed a slightly smaller mean EOO (no difference in specimen count) than terrestrial species, while epiphytic Ericaceae exhibited larger EOO and specimen count than terrestrial species.

**Figure 3:**
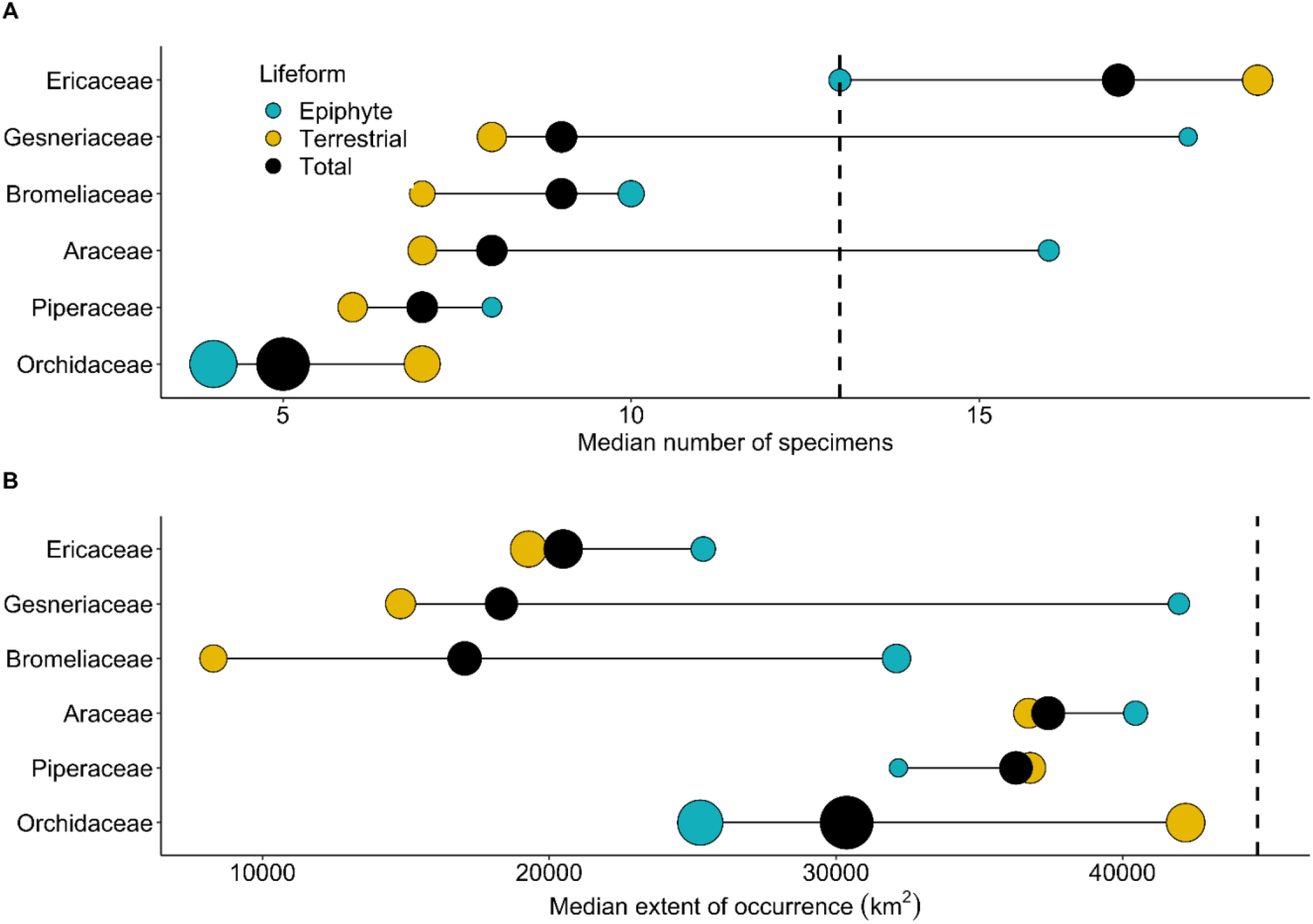
The median range size of epiphyte-rich families is typically smaller than the angiosperm-wide median range size. Bubbles show median range size measured as A) specimen count and B) extent of occurrence (EOO) of epiphytic, terrestrial, and all species for each of the six most epiphyte-rich families. The median range size of angiosperms (EOO=44,326 km^2^, specimen count=13) is shown as a vertical dotted line. Bubble size is proportional to the number of species in each group.

Species in epiphyte-rich families (including both epiphytic and terrestrial species) have, on average, 53% (95% CI: 52%-54%) fewer specimens and 42% (95% CI: 40%-44%) smaller EOOs than species in other angiosperm families. The median specimen count in five of the six most epiphyte-rich families was smaller than the angiosperm-wide average (excepting Ericaceae, Fig. 3). All epiphyte-rich families had smaller median EOOs than the angiosperm-wide median. These results suggest that belonging to an epiphyte-rich family has a similar or stronger effect on range size than being epiphytic.

### Analyses controlling for evolutionary relatedness

Controlling for covariation due to shared ancestry revealed that the tendency for angiosperm epiphytes to have small ranges is not caused by epiphytism. In fact, in a phylogenetic generalized least squares (PGLS) regression model, epiphytes had significantly larger range sizes than terrestrial plants: on average 46% (95% CI: 43%-49%) more specimens and 54% (95% CI: 47%-61%) larger EOOs (Fig. 2), suggesting that epiphytism per se contributes to larger range sizes. Values are mean estimates from regressions using 100 phylogenetic trees with missing species randomly imputed, which provides an estimate of the uncertainty associated with the imputation approach (see Tables S1-S2).

The phylogenetically independent effects of epiphytism were associated with considerably larger mean EOOs and larger mean specimen counts in three epiphyte-rich families: Bromeliaceae, Araceae, and Gesneriaceae (Fig. 2). The remaining families showed either no effects of epiphytism on mean EOO and a positive effect on specimen counts (Orchidaceae and Piperaceae) or no effect (Ericaceae). However, in tropical-only analyses, epiphytic Ericaceae showed larger EOO and specimen count, exhibiting a similar pattern to the primarily tropical families Bromeliaceae and Gesneriaceae.

As in ordinary linear regressions, range size comparison between lifeforms within families showed that epiphytes were more likely to differ from terrestrial species in their mean specimen count (67% of families) than their EOO (17% of families). In phylogenetic models, epiphytism was not associated with a decrease in mean EOO or specimen count for any of the 24 families with 10 or more epiphytic species, except for the small Neotropical family Schlegeliaceae, indicating that epiphytism is rarely functionally associated with range sizes smaller than those of confamilial terrestrial species (Tables S1 and S2).

### Conservation implications

Of the 8,170 epiphyte species for which EOO was estimated, 45% had EOO <20,000 km^2^, the threshold value for categorisation as Vulnerable following Red List Criterion B1^32^. These species can be considered to have ‘vulnerably small ranges’^22^ but would need to meet additional subcriteria to be categorised as Vulnerable in a Red List assessment^32^. Of the 20,326 epiphyte species for which specimen count was estimated, 51% had ≤5 specimens and are considered ‘rare’^33^. In contrast, 39% of the 137,728 terrestrial species with EOO estimates had vulnerably small ranges, and 29% of the 224,135 terrestrial species were considered rare, showing that epiphytes had a consistently higher proportion of species with vulnerably small range sizes compared to terrestrial species. The precise global proportion of species with vulnerably small ranges is still unknown because 27% of angiosperms were not included in this analysis, due to their lack of digitally available specimen records (11%) or their occurrence in five or more botanical countries (16%).

## Discussion

Analyses controlling for phylogenetic relatedness reveal that epiphytes are associated with larger range sizes than closely related terrestrial plant species. By comparing phylogenetically controlled and uncontrolled regressions, we infer the effect of evolution on the epiphytism-range size relationship and conclude that the observed small ranges of epiphytes are the consequence of their concentration in a few species-rich lineages with generally small-ranged species. In other words, epiphytism does not cause small ranges. Over 90% of angiosperm epiphyte diversity is concentrated in just six families, indicating that relatively few lineages have effectively radiated in the epiphytic niche. All these epiphyte-rich families (except Ericaceae) have median specimen counts and EOOs smaller than the angiosperm-wide values (Fig. 3) and their species have smaller mean specimen counts and EOOs than other angiosperm families.

Traits predisposing species to small range sizes may be prevalent in epiphyte-rich clades. The epiphytic habitat is characterised by a set of key challenges, including limited and variable water supplies^34^, low substrate stability^35^, and discontinuous habitat^36^. Epiphytes possess unique trait syndromes compared to ground-rooted plants^37^, and diverse morphological and physiological traits observed in epiphytes have been interpreted as adaptations for arboreal life^4^. However, traits advantageous in the epiphytic habitat are often shared between epiphytes and closely related terrestrial species, suggesting that they are not necessarily adaptations for epiphytism. For instance, structures explained in terms of desiccation avoidance or tolerance in epiphytic taxa are often found across entire clades containing epiphytes. The absorptive velamen radicum root epidermis, typically considered an adaptation in epiphytic orchids and some aroids, is found throughout monocotyledons, including over 160 terrestrial orchid genera^38^. The water-impounding tank structures (‘phytotelmata’) and absorptive leaf scales of bromeliads characterise both terrestrial and epiphytic taxa within specific subclades^39^. The abundance of taxa exhibiting various degrees of facultative epiphytism^5^ and the apparently numerous gains and losses of epiphytism in epiphyte-rich clades such as orchids also indicate that epiphytism is an evolutionarily labile trait within certain lineages^7,40^.

Epiphytic and terrestrial taxa within certain clades are not only morphologically similar, but also ecologically similar. Lithophytes, plants which grow on rocks (here included in terrestrial plants), face challenges associated with growth on largely impenetrable substrates, like those faced by epiphytes. The lithophytic lifeform is common in epiphyte-rich families, including Orchidaceae (e.g. *Epidendrum*, *Laelia*, *Cyrtopodium*), Bromeliaceae (e.g. *Encholirium*, *Pitcairnia*, *Tillandsia*), Araceae (e.g. *Anthurium*), and Gesneriaceae (e.g. *Sinningia*)^15,41^. Many species in these families grow as either epiphytes or lithophytes^4^, exemplifying the similarity of these habitats and their demands. Thus, epiphyte-rich families may possess traits adapted for stressful conditions including, but not limited to, the epiphytic habitat. The tendency for entire epiphyte-rich families to have small ranges may be linked to their specialism in stressful habitats. Niche breadth is positively correlated with range size^42^, and other groups of substrate-specialist plant species such as those restricted to rocky outcrops^1^ and serpentine soils^43^ also tend toward small range sizes.

While epiphytism itself may not be functionally related to small ranges, evolution of key traits underpinning colonisation of and radiation within the epiphytic habitat may indirectly yield the small ranges observed in epiphyte-rich clades by triggering speciation and increasing net diversification. Epiphyte-rich clades are notably speciose at multiple taxonomic scales: Orchidaceae (69% epiphytic) is the most speciose plant family and Bromeliaceae (55% epiphytic) is among the most species-rich in the Neotropics. Epiphyte-dominated genera are among the most species-rich in plants, e.g. *Bulbophyllum* (Orchidaceae), 2,106 species; *Peperomia* (Piperaceae), 1,409 species; *Anthurium* (Araceae), 1,137 species^30^. Studies integrating phylogenetic and trait data link the remarkable species diversity of orchids and bromeliads to accelerations in diversification exceeding average rates for angiosperms. In Bromeliaceae, evolution of phytotelmata, wind- and bird-dispersed seeds, and epiphytism are linked to accelerated diversification^44^. In orchids, diversification is linked to evolution of pollinia, CAM photosynthesis, and epiphytism^45^. Both families diversified in tropical cordilleras characterised by topographical complexity and barriers to gene flow, perhaps fostering genetic isolation of small populations, begetting many closely related, small-ranged species^44,45^.

In analyses controlling for effects of shared ancestry, epiphytes are associated with larger ranges than terrestrial species across angiosperms and within most epiphyte-rich families. Dispersal ability is positively associated with range size in vascular plants, albeit variably between clades^19^. Epiphyte-rich lineages are characterised by seed-types well-adapted for tree-to-tree dispersal, including light, wind-dispersed seeds (e.g. dust-like seeds in orchids, plumed seeds in Tillandsioideae bromeliads and Gesneriaceae) and fleshy, bird- or bat-dispersed seeds (e.g. Bromelioideae bromeliads; *Anthurium* (Araceae); *Piper* (Piperaceae)^4,7,13,46,47^. Although seed weight in epiphytes follows a bimodal distribution (consistent with the hypothesis that light wind-dispersed and heavier, fleshy bird-dispersed seeds are both effective for tree-to-tree dispersal), epiphytes do not generally have smaller seeds than confamilial terrestrial plants, evidence that closely related epiphytic and terrestrial taxa exhibit similar traits^47^. However, dispersal of anemochorous seeds is likely more successful from greater heights in the canopy than from the ground, thereby predisposing epiphytic taxa to larger ranges than morphologically-similar, closely related terrestrial taxa^17^. Bird-dispersed epiphyte seeds may also reach greater distances if canopy-inhabiting birds and bats forage over greater distances than forest-floor birds^14,44^.

In certain families, the much larger ranges of epiphytes compared to terrestrial species could be partly attributable to the exceptionally small ranges of their terrestrial species. Notably, terrestrial bromeliads and gesneriads have median EOOs 5 and 3 times smaller than the angiosperm-wide median, respectively (Fig. 3). Terrestrial species endemism is frequently driven by substrate specialisation^48^, whereas epiphytes generally show little preference for particular host tree species^49^. Substrate-driven endemism may be particularly marked in lithophytes occurring on highly-fragmented and isolated island-like rock outcrops such as inselbergs^41^. In Brazil and Madagascar, centres of diversity for epiphytes^6^, inselberg floras are exceptionally rich in endemic species^41^. Thus, the small ranges of some terrestrial species in epiphyte-rich families, including bromeliads and gesneriads, may be driven by lithophytes and other endemics of isolated and fragmented habitats.

The substantial correlation (0.6) between EOO and specimen count supports the notion that the number of herbarium records in GBIF reflects differences in species’ geographic range size. The direction of the effect of epiphytism was consistent between the two metrics across 24 epiphytic families. However, in ten families, epiphytes had greater numbers of specimens yet showed no difference in their EOO. Epiphytes are better represented in our specimen count dataset (8.4% of species) than in the EOO dataset (5.6% of species, Table S6) because epiphytes are generally so rare or poorly-known that a disproportionate number of them lack the three georeferenced occurrence records required for EOO estimation. Thus, the specimen count metric may be more likely to detect differences in range size between epiphytes and terrestrial plants because it includes the rarest epiphytic species. Patterns of habitat availability for epiphytes may differ from those of soil-rooted plants: niche differentiation can be high on a single host tree^50^ but low between different hosts^51^. Thus, local epiphyte quotients can be extremely high^8^ but decrease with area as different terrestrial niches, and their species, are captured^51^. Lithophytes and other substrate-specialist terrestrial species may have highly fragmented distributions within their ranges if they are restricted to small, patchily distributed outcrops^41,48^.

Phylogenetic regression models are methods of choice in studying relationships between species’ traits^52^. We used them to show that epiphytism is functionally associated with larger range sizes in angiosperms despite epiphytes having significantly smaller average range sizes than the angiosperm average. However, small-ranged species are vulnerable to extinction regardless of the drivers of their range size, and species-level conservation prioritisation typically treats species as independent units. Contrary to recommendations that raw-data analyses are unnecessary when phylogenetic data is available^52^, we re-emphasise that ordinary, non-phylogenetic models are useful to uncover correlates of extinction risk and, in conjunction with phylogenetic models, infer evolutionary effects on relationships between range size and species traits^1^. Although epiphytism explains relatively little variation in range size for angiosperms, combined effects of multiple traits can achieve substantial predictive power in range size analyses^53^. Identifying correlates of rarity informs conservation prioritisation, for instance, as explanatory variables in machine-learning models used to predict extinction risk^54,55^.

Angiosperm epiphytes are most diverse and have their smallest ranges in tropical regions which are experiencing some of the highest rates of habitat loss and degradation globally^56^. Epiphytes may be particularly vulnerable to extinction^54^, even when controlling for range size^1^. However, in Brazil’s Atlantic rainforest, lifeform predicts extinction risk only in non-phylogenetic models^1^, suggesting that vulnerability may be a characteristic of entire epiphyte-rich lineages^57^, consistent with the relationship between epiphytism and range size demonstrated here. Many epiphytes and their close relatives exhibit slow life-histories, predisposing them to slow population recovery following disturbances^58–60^. Species in epiphyte-rich clades are also particularly threatened by overcollection for horticultural use^61^, a threat potentially exacerbated by slow population recovery.

The arboreal habitat of epiphytes may be uniquely impacted by certain anthropogenic activities. Epiphyte diversity is lower in secondary and fragmented forests than primary contiguous forests^62,63^. Fragmentation and degradation of tropical forests alter tree canopy microclimates with direct impacts on epiphyte persistence^64^ and indirect impacts including increase in lianas which may outcompete epiphytes^65^. Selective logging of large, old trees may remove hosts with the most diverse epiphyte assemblages due to their more varied microclimates and longer time for colonisation^50,62^.

Despite their unusual ecology, inherently small ranges, and apparent heightened vulnerability to extinction, the conservation status of most epiphytes remains unknown. Our findings that 45% of epiphytes have vulnerably small EOOs and that 51% qualify as rare, are cause for grave concern for the survival of these species, a concern amplified by the fact that only 7% of epiphytic angiosperms have Red List assessments^66^. Some of the lineages identified here as having particularly small ranges, such as bromeliads and epiphytic orchids, are particularly poorly represented on the Red List^67,68^. This is especially troubling considering that tropical plants and plant groups wild-harvested for horticulture are reported to have especially high proportions of threatened plants^69,70^. Thus, Red Listing of epiphytes and epiphyte-rich lineages should be prioritised to enhance understanding of the threats they face and inform conservation action for the high proportion likely threatened with extinction.

## Methods

### Data

The World Checklist of Vascular Plants (WCVP) is the first comprehensive, continually updated source for vascular plant species names and geographic distributions^30^. We used WCVP to obtain taxonomic and geographic distribution data for all 336,281 accepted angiosperm species not of hybrid origin and with known distributions. We intersected WCVP and EpiList 1.0, a comprehensive list of epiphytes, to categorise over 27,000 angiosperm species as epiphytes for this analysis^5^. Hemiepiphytes, which germinate epiphytically but later root in the ground, were excluded from this analysis because their ecophysiology differs from true epiphytes^71^. Restricting our analysis to angiosperms prevented over-complicating inferences concerning determinants of range size by excluding pteridophytes, which have diverging physiology and distribution patterns^4,6^.

We calculated three species-level range size metrics from three different types of occurrence data: number of botanical countries with native occurrences, extent of occurrence (EOO) calculated from georeferenced herbarium data, and total number of herbarium specimens including both georeferenced and non-georeferenced records. ‘Botanical countries’ is the standard unit for recording plant distributions in WCVP following the World Geographical Scheme for Recording Plant Distributions (WGSRPD)^29^. Globally, the 368 botanical countries mostly correspond to political countries, although some large countries are split into smaller units, and geographically disjunct areas (e.g. islands) may be recognised as distinct botanical countries, reflecting their phytogeographical differences^29^.

We used WCVP’s number of botanical countries to estimate range size for all vascular plant species. The number of botanical countries captures the variation in EOO remarkably well among the large-ranged species and probably around half of the total variation in EOO^54^. However, the coarse resolution of botanical countries misses range-size variation among small-ranged species. Species occurring in four or fewer botanical countries can meet at a single point^29^; thus, narrow-ranged endemics with extremely small EOOs could have native occurrences in up to four botanical countries. To account for variation among rare species, we calculate two additional metrics of range size for angiosperms occurring in ≤4 botanical countries (84% of all angiosperm species): sum of unique herbarium specimen records for a species (hereafter ‘specimen count’) and EOO. Occurrence records from preserved specimens were obtained from the Global Biodiversity Information Facility (GBIF.org; see Table S5 for download citations) using the R package ‘rgbif’^72^. Only herbarium specimen-based occurrence records were used for our calculations because they represent verifiable evidence of species occurrences and underpin accurate estimations of species distributions even at relatively low sampling densities^73,74^.

### Data cleaning

Spatial and taxonomic biases, coordinate inaccuracies, and duplication of records are well-documented issues with aggregated biodiversity databases, including GBIF^75,76^. To mitigate these issues, we employed two different data cleaning procedures for georeferenced records and non-georeferenced records, respectively. Among georeferenced occurrence records, we detected records of the same species with identical coordinates (rounded to three decimal places), removed duplicates and retained at most one record per 110 m^2^ for each species. We used the R package ‘rWCVP’ to remove records outside each species’ reported native range at botanical country level^77^. We removed records with low precision (coordinate uncertainty >100 km). Finally, automatic filters from the R package ‘CoordinateCleaner’ were applied to remove potentially erroneous or unfit records^78^. Records with latitude = longitude, or both = 0 were removed, as were records for which coordinates corresponded to country centroids or capitals, or biodiversity institutions.

To clean non-georeferenced occurrence records, for each species we removed records with duplicated collection locality or duplicated year and administrative unit (‘stateProvince’ in GBIF) as these likely represent duplicates of the same population or collection event, respectively. Non-georeferenced records with species’ native countries of occurrence were removed by mapping botanical countries to ISO country codes in GBIF records and removing records from outside the species’ native ranges as recorded in WCVP. In total, 11,549,160 occurrence records were retained after data-cleaning, of which 5,934,797 were georeferenced and 5,614,363 were non-georeferenced.

### Range sizes and phylogeny

The number of native, occupied botanical countries for 336,281 accepted angiosperm species were obtained directly from WCVP. The EOO was estimated using cleaned georeferenced occurrence records for species having ≥3 georeferenced records. Using the R package rCAT, we calculated EOO as the minimum convex polygon encompassing all occurrence points for a species^79^.

We used specimen count as a complementary measure of range size and a correlate of the global abundance and occupied area of each species, rather than area of occupancy (AOO). This helped maximise the number of species included in our study, as 48.6% of the cleaned GBIF records are not georeferenced, and 58,974 angiosperm species occurring in ≤4 botanical countries had only non-georeferenced GBIF records. Omitting non-georeferenced occurrence data risks underestimating species range size^73^ and amplifying biases against species lacking georeferenced records which are likely to be poorly known and rare. We used cleaned georeferenced and non-georeferenced occurrence records to calculate specimen counts. Since AOO is often calculated as the number of 2×2 km^2^ grid cells occupied by a species^32^ and, after cleaning, each herbarium specimen theoretically represents a unique locality, AOO and specimen count should be highly correlated, and studies using the number of specimens as a correlate of global abundance of individual species^33^ and rarity^73^ report good performance against expert-backed assessments.

To control for phylogenetic distances between species spanning the angiosperm clade, we used a set of 100 species-level phylogenetic trees for angiosperms prepared with 329,798 accepted angiosperms^31^. This tree uses Smith & Brown’s GBMB phylogenetic tree^80^ as a backbone, updated to reflect recent taxonomic changes to angiosperm families, genera, and species as published in WCVP (version 6). Missing genera and species were randomly added at the appropriate crown nodes, producing 100 trees by repeating the imputation stage 100 times to account for uncertainty in phylogenetic placement of these taxa. All names in the phylogenetic trees are standardised to WCVP, facilitating name-matching between range size data and tree tip labels for phylogenetic regression.

Our dataset contained specimen counts for 244,461 species (96.5% of species with specimen counts) and EOOs for 145,898 species (99.2% of species with EOO). The difference in species numbers between the phylogenetic trees and the distribution data is due to nomenclatural and taxonomic changes in the WCVP which took place between the version of the WCVP used to build the trees (version 6) and the Special Issue version^81^ used for this analysis.

After data cleaning and name-matching with a phylogenetic tree, we achieved 100% coverage of angiosperm species with one range size metric (number of botanical countries; used only for ordinary regression), 72.7% coverage of angiosperms and 73.8% of epiphytes with two range size metrics (botanical countries and specimen count) and 43.4% coverage of angiosperms and 29.6% of epiphytes with all three range size metrics.

The percentage of epiphytes for which EOO could be calculated varied across the six most epiphyte-rich families, being the lowest in Orchidaceae (25%) and highest in Araceae (72%). Most of this variation was due to species, especially orchids, having <3 georeferenced records, preventing calculation of EOO. The percentage of epiphytes for which specimen counts were available was more consistent across families (Table S6). Across epiphyte-rich families, coverage by at least one GBIF-derived metric was lowest for Orchidaceae (78%) and highest for Bromeliaceae (92%). We generated correlation plots and calculated correlation coefficients between each pair of the three range size metrics (Fig. S1).

### Statistical analyses

Regression methods were used to test the relationships between lifeform (epiphyte vs terrestrial) and each of the three range size metrics across angiosperms. We also investigated differences in range size and the effect of epiphytism within each of the twenty-four families with ≥10 epiphytic species (Table S4) using each of the three metrics (Tables S1-3). All statistical analyses were conducted in the R environment^81^ (version 4.1.2). We conducted a sensitivity analysis restricted to angiosperms for which the mean latitude of their distribution, measured as botanical countries, is in the tropics (n=195,312; Fig. S2) to test whether our results are confounded by the largely tropical distribution of epiphytes^6^ and/or the tendency for range size to increase with latitude^82^. A further sensitivity analysis excluded 19,915 angiosperm species occurring on oceanic islands (n=316,366; Fig. S3), which have different biogeographical and evolutionary histories than continental islands and mainland regions^83^.

All three range size metrics were highly positively skewed due to the large proportion of rare species. We used generalised linear regression with a quasipoisson model to test relationships between the number of botanical countries and lifeform (epiphyte or terrestrial). For both specimen count and EOO, we applied the Box-Cox method^84^ and identified the logarithmic scale as the most suitable transformation for meeting the assumption of symmetry in the data for linear regression. For specimen count and EOO metrics, a phylogenetic regression was implemented in addition to ordinary linear regression. Phylogenetic regression estimates the independent effect of predictors on range size by controlling for the non-independence between species caused by shared ancestry. Since closely related species are more likely to have similar range sizes than expected by chance^26^, phylogenetic regression is more appropriate for testing the causality of epiphytism on range size than methods which do not control for shared ancestry^52^. Standard linear regression allows the comparison of unweighted range size values, offering useful context for comparison and information for conservation prioritisation, which typically treats species as independent units. We used a quasipoisson model to account for overdispersion in the relationship between number of botanical countries and lifeform, due to low variation and prevalence of species found in a single botanical country (65%). We could not satisfactorily employ the phylogenetic regression method using number of botanical countries.

We used the function ‘phylolm’ from the R package ‘phylolm’ to carry out phylogenetic regression^85^. ‘Phylolm’ uses the phylogenetic generalised least squares (PGLS) method to correct the error structure of the residuals of the linear model to account for covariation due to shared ancestry. We used Pagel’s *λ* to estimate the strength of the phylogenetic signal in the model’s residuals. Pagel’s *λ* is derived through maximum likelihood estimation and is multiplied by the expected variance-covariance matrix to account for the strength of the phylogenetic signal in the data^86^. Values of *λ* approaching 1 indicate very strong phylogenetic correlation between species while values of *λ* close to 0 indicate weak phylogenetic correlation. The ‘phylolm’ function calculates *λ* based on the residual errors of the model and does not reflect the phylogenetic signal of the response variable itself, but rather of the relationship between the predictor(s) and response variable. Range size heritability varies between groups, and phylogenetic signal in the relationship between lifeform and range size is unlikely to be fixed across angiosperms: Pagel’s *λ* accounts for this variability. Phylogenetic regression models were run for the entire angiosperm dataset using each of the 100 phylogenetic trees^31^. The mean and standard error of parameter estimates were calculated to account for uncertainty in the phylogenetic placement of certain taxa^89^.

To investigate how the range size of epiphyte-rich groups compares to other angiosperm lineages, we tested if the mean specimen count and EOO for species in each of the six most epiphyte-rich families differed from that of species in other families. The most epiphyte-rich families were Orchidaceae (n=21,034 epiphyte species), Bromeliaceae (n=1,937), Ericaceae (n=876), Araceae (n=766), Piperaceae (n=699), and Gesneriaceae (n=644). These families include >90% of epiphyte species and represent lineages which have effectively radiated in the epiphyte niche^7^. For each family, we plotted the percentage difference between mean epiphyte range sizes and terrestrial plant range size across all three metrics for both phylogenetic and ordinary regression models, to facilitate cross-metric comparison of the effect of epiphytism on range size in these lineages (Fig. 2).

To explore our findings in a conservation context, we calculated the proportion of epiphyte species with EOO <20,000 km^2^, the threshold for consideration of species as threatened under Red List Criterion B1^32^, though further subcriteria must be considered before a species is categorised. We also calculated the proportion of epiphyte species with specimen count ≤ 5, which may be considered rare following^33^ Enquist *et al*. (2019).

## Data availability

We confirm that if the manuscript is accepted, the data supporting the results will be archived in an appropriate public repository and the data DOI will be included at the end of the article.

## Acknowledgements

We thank Matilda Brown for their suggestions and support regarding statistics and figures.

## Author contributions

TCCL and ENL conceived the initial idea. TCCL, ENL, and VJS designed the research. FF compiled and provided the phylogenetic trees for statistical analyses. VJS compiled, analysed and interpreted the data. VJS wrote the initial draft of the manuscript with input from ENL, TCCL, and FF. All authors contributed to the final version of this paper.

## References

1. Leão, T. C. C., Fonseca, C. R., Peres, C. A. & Tabarelli, M. Predicting Extinction Risk of Brazilian Atlantic Forest Angiosperms. Conserv. Biol. 28, 1349–1359 (2014).

2. Gaston, K. J. & Fuller, R. A. The sizes of species’ geographic ranges. J. Appl. Ecol. 46, 1–9 (2009).

3. Keith, D. A., Akçakaya, H. R. & Murray, N. J. Scaling range sizes to threats for robust predictions of risks to biodiversity. Conserv. Biol. 32, 322–332 (2018).

4. Zotz, G. Plants on Plants – The Biology of Vascular Epiphytes. (Springer International Publishing, 2016). doi:10.1007/978-3-319-39237-0.

5. Zotz, G., Weigelt, P., Kessler, M., Kreft, H. & Taylor, A. EpiList 1.0: a global checklist of vascular epiphytes. Ecology 102, e03326 (2021).

6. Taylor, A. et al. Vascular epiphytes contribute disproportionately to global centres of plant diversity. Glob. Ecol. Biogeogr. 31, 62–74 (2022).

7. Gentry, A. H. & Dodson, C. H. Diversity and Biogeography of Neotropical Vascular Epiphytes. Ann. Mo. Bot. Gard. 74, 205–233 (1987).

8. Kelly, D. L., Tanner, E. V. J., Lughadha, E. M. N. & Kapos, V. Floristics and Biogeography of a Rain Forest in the Venezuelan Andes. J. Biogeogr. 21, 421–440 (1994).

9. Stuntz, S., Ziegler, C., Simon, U. & Zotz, G. Diversity and structure of the arthropod fauna within three canopy epiphyte species in central Panama. J. Trop. Ecol. 18, 161–176 (2002).

10. Mccracken, S. F. & Forstner, M. R. J. Herpetofaunal community of a high canopy tank bromeliad (Aechmea zebrina) in the Yasuní Biosphere Reserve of Amazonian Ecuador, with comments on the use of “arboreal” in the herpetological literature. Amphib. Reptile Conserv. 8, 65–75 (2014).

11. Gotsch, S. G., Nadkarni, N. & Amici, A. The functional roles of epiphytes and arboreal soils in tropical montane cloud forests. J. Trop. Ecol. 32, 455–468 (2016).

12. Schimper, A. F. W. Die epiphytische Vegetation Amerikas. (Fischer, 1888).

13. Zotz, G. Biogeography: Latitudinal and Elevational Trends. in Plants on Plants – The Biology of Vascular Epiphytes (ed. Zotz, G.) 51–66 (Springer International Publishing, 2016). doi:10.1007/978-3-319-39237-0_3.

14. Ibisch, P. L., Boegner, Andreas, Nieder, Jürgen & Barthlott, W. How diverse are neotropical epiphytes? An analysis based on the ‘Catalogue of the flowering plants and gymnosperms of Peru’. Ecotropica 2, 13–28 (1996).

15. Kessler, M. Environmental patterns and ecological correlates of range size among bromeliad communities of Andean Forests in Bolivia. Bot. Rev. 68, 100–127 (2002).

16. Martins, A. C. et al. From tree tops to the ground: Reversals to terrestrial habit in Galeandra orchids (Epidendroideae: Catasetinae). Mol. Phylogenet. Evol. 127, 952–960 (2018).

17. Reimuth, J. & Zotz, G. The biogeography of the megadiverse genus Anthurium (Araceae). Bot. J. Linn. Soc. 194, 164–176 (2020).

18. Leão, T., Reich, P. & Lughadha, E. N. The geographic range size and vulnerability to extinction of epiphytes in the Atlantic Forest of Brazil. https://www.authorea.com/users/500354/articles/581156-the-geographic-range-size-and-vulnerability-to-extinction-of-epiphytes-in-the-atlantic-forest-of-brazil?commit=d7f3f69ff81e7f452a7f27360895134c6cf48aa5 (2022) doi:10.22541/au.166012202.29914221/v1.

19. Alzate, A. & Onstein, R. E. Understanding the relationship between dispersal and range size. Ecol. Lett. 25, 2303–2323 (2022).

20. Katul, G. G. et al. Mechanistic Analytical Models for Long-Distance Seed Dispersal by Wind. Am. Nat. 166, 368–381 (2005).

21. Davies, T. J. et al. Extinction Risk and Diversification Are Linked in a Plant Biodiversity Hotspot. PLOS Biol. 9, e1000620 (2011).

22. Leão, T. C. C., Lughadha, E. N. & Reich, P. B. Evolutionary patterns in the geographic range size of Atlantic Forest plants. Ecography 43, 1510–1520 (2020).

23. Testo, W. L., Sessa, E. & Barrington, D. S. The rise of the Andes promoted rapid diversification in Neotropical *Phlegmariurus* (Lycopodiaceae). New Phytol. 222, 604–613 (2019).

24. Kreft, H., Köster, N., Küper, W., Nieder, J. & Barthlott, W. Diversity and biogeography of vascular epiphytes in Western Amazonia, Yasuní, Ecuador. J. Biogeogr. 31, 1463–1476 (2004).

25. Jablonski, D. Heritability at the Species Level: Analysis of Geographic Ranges of Cretaceous Mollusks. Science 238, 360–363 (1987).

26. Hunt, G., Roy, K. & Jablonski, D. Species-Level Heritability Reaffirmed: A Comment on “On the Heritability of Geographic Range Sizes.” Am. Nat. 166, 129–135 (2005).

27. Borregaard, M. K., Gotelli, N. J. & Rahbek, C. ARE RANGE-SIZE DISTRIBUTIONS CONSISTENT WITH SPECIES-LEVEL HERITABILITY? Evolution 66, 2216–2226 (2012).

28. Freckleton, R. P., Harvey, P. H. & Pagel, M. Phylogenetic Analysis and Comparative Data: A Test and Review of Evidence. Am. Nat. 160, 712–726 (2002).

29. Brummit, R.K. World geographical scheme for recording plant distributions, edn 2. Plant taxonomic database standards no. 2. edn. 2. (Hunt Institute for Botanical Documentation, 2001).

30. Govaerts, R., Nic Lughadha, E., Black, N., Turner, R. & Paton, A. The World Checklist of Vascular Plants, a continuously updated resource for exploring global plant diversity. Sci. Data 8, 215 (2021).

31. Forest, F. Species-level phylogenetic trees of all angiosperm species (100 trees). (2023) 10.5281/zenodo.7600341.

32. IUCN Red List categories and criteria, version 3.1*, second edition*. (IUCN, 2012).

33. Enquist, B. J. et al. The commonness of rarity: Global and future distribution of rarity across land plants. Sci. Adv. 5, eaaz0414 (2019).

34. Zotz, G. & Hietz, P. The physiological ecology of vascular epiphytes: current knowledge, open questions. J. Exp. Bot. 52, 2067–2078 (2001).

35. Cabral, J. S. et al. Branchfall as a Demographic Filter for Epiphyte Communities: Lessons from Forest Floor-Based Sampling. PLOS ONE 10, e0128019 (2015).

36. Laube, S. & Zotz, G. A metapopulation approach to the analysis of long-term changes in the epiphyte vegetation on the host tree Annona glabra. J. Veg. Sci. 18, 613–624 (2007).

37. Hietz, P. et al. Putting vascular epiphytes on the traits map. J. Ecol. 110, 340–358 (2022).

38. Zotz, G., Schickenberg, N. & Albach, D. The velamen radicum is common among terrestrial monocotyledons. Ann. Bot. 120, 625–632 (2017).

39. Benzing, D. H. Bromeliaceae: Profile of an Adaptive Radiation. (Cambridge University Press, 2000).

40. Chomicki, G. et al. The velamen protects photosynthetic orchid roots against UV-B damage, and a large dated phylogeny implies multiple gains and losses of this function during the Cenozoic. New Phytol. 205, 1330–1341 (2015).

41. Porembski, S. Tropical inselbergs: habitat types, adaptive strategies and diversity patterns. Rev. Bras. Botânica 30, 579–586 (2007).

42. Kambach, S. et al. Of niches and distributions: range size increases with niche breadth both globally and regionally but regional estimates poorly relate to global estimates. Ecography 42, 467–477 (2019).

43. Essl, F. et al. Distribution patterns, range size and niche breadth of Austrian endemic plants. Biol. Conserv. 142, 2547–2558 (2009).

44. Givnish, T. J. et al. Adaptive radiation, correlated and contingent evolution, and net species diversification in Bromeliaceae. Mol. Phylogenet. Evol. 71, 55–78 (2014).

45. Givnish, T. J. et al. Orchid phylogenomics and multiple drivers of their extraordinary diversification. Proc. R. Soc. B Biol. Sci. 282, 20151553 (2015).

46. Salazar, D., Kelm, D. H. & Marquis, R. J. Directed seed dispersal of Piper by Carollia perspicillata and its effect on understory plant diversity and folivory. Ecology 94, 2444–2453 (2013).

47. Rockwood, L. L. Seed Weight as a Function of Life Form, Elevation and Life Zone in Neotropical Forests. Biotropica 17, 32–39 (1985).

48. Morat, Ph. Our Knowledge of the Flora of New Caledonia: Endemism and Diversity in Relation to Vegetation Types and Substrates. Biodivers. Lett. 1, 72–81 (1993).

49. Wagner, K., Mendieta-Leiva, G. & Zotz, G. Host specificity in vascular epiphytes: a review of methodology, empirical evidence and potential mechanisms. AoB PLANTS 7, plu092 (2015).

50. Woods, C. L., Cardelús, C. L. & DeWalt, S. J. Microhabitat associations of vascular epiphytes in a wet tropical forest canopy. J. Ecol. 103, 421–430 (2015).

51. Nieder, J., Engwald, S. & Barthlott, W. Patterns of Neotropical Epiphyte Diversity. Selbyana 20, 66–75 (1999).

52. Symonds, M. R. E. & Blomberg, S. P. A Primer on Phylogenetic Generalised Least Squares. in Modern Phylogenetic Comparative Methods and Their Application in Evolutionary Biology: Concepts and Practice (ed. Garamszegi, L. Z.) 105–130 (Springer, 2014). doi:10.1007/978-3-662-43550-2_5.

53. Cardillo, M. & Skeels, A. Spatial, Phylogenetic, Environmental and Biological Components of Variation in Extinction Risk: A Case Study Using Banksia. PLOS ONE 11, e0154431 (2016).

54. Bachman, S. P., Brown, M. J. M., Leão, T. C. C., Lughadha, E. N. & Walker, B. E. Extinction risk predictions for the world’s flowering plants to support their conservation. 2023.08.29.555324 Preprint at 10.1101/2023.08.29.555324 (2023).

55. Walker, B. E., Leão, T. C. C., Bachman, S. P., Lucas, E. & Nic Lughadha, E. Evidence-based guidelines for automated conservation assessments of plant species. Conserv. Biol. 37, e13992 (2023).

56. Hansen, M. C. et al. High-Resolution Global Maps of 21st-Century Forest Cover Change. Science 342, 850–853 (2013).

57. Leão, T. C. C., Reich, P. B. & Nic Lughadha, E. The geographic range size and vulnerability to extinction of angiosperm epiphytes in the Atlantic Forest of Brazil. Glob. Ecol. Biogeogr. **n/a**,.

58. Zotz, G. How Fast Does an Epiphyte Grow? Selbyana 16, 150–154 (1995).

59. Schmidt, G. & Zotz, G. Inherently slow growth in two Caribbean epiphytic species: A demographic approach. J. Veg. Sci. 13, 527–534 (2002).

60. Einzmann, H. J. R., Weichgrebe, L. & Zotz, G. Long-term community dynamics in vascular epiphytes on Annona glabra along the shoreline of Barro Colorado Island, Panama. J. Ecol. 109, 1931–1946 (2021).

61. Flores-Palacios, A. & Valencia-Díaz, S. Local illegal trade reveals unknown diversity and involves a high species richness of wild vascular epiphytes. Biol. Conserv. 136, 372–387 (2007).

62. Köster, N., Nieder, J. & Barthlott, W. Effect of Host Tree Traits on Epiphyte Diversity in Natural and Anthropogenic Habitats in Ecuador. Biotropica 43, 685–694 (2011).

63. Barthlott, W., Schmit-Neuerburg, V., Nieder, J. & Engwald, S. Diversity and abundance of vascular epiphytes: a comparison of secondary vegetation and primary montane rain forest in the Venezuelan Andes. Plant Ecol. 152, 145–156 (2001).

64. Werner, F. A. Reduced growth and survival of vascular epiphytes on isolated remnant trees in a recent tropical montane forest clear-cut. Basic Appl. Ecol. 12, 172–181 (2011).

65. Campbell, M. J. et al. Edge disturbance drives liana abundance increase and alteration of liana–host tree interactions in tropical forest fragments. Ecol. Evol. 8, 4237–4251 (2018).

66. The IUCN Red List of Threatened Species. IUCN Red List of Threatened Species https://www.iucnredlist.org/en.

67. Zizka, A. et al. Biogeography and conservation status of the pineapple family (Bromeliaceae). Divers. Distrib. 26, 183–195 (2020).

68. Zizka, A., Silvestro, D., Vitt, P. & Knight, T. M. Automated conservation assessment of the orchid family with deep learning. Conserv. Biol. 35, 897–908 (2021).

69. Brummitt, N. A. et al. Green Plants in the Red: A Baseline Global Assessment for the IUCN Sampled Red List Index for Plants. PLOS ONE 10, e0135152 (2015).

70. Goettsch, B. et al. High proportion of cactus species threatened with extinction. Nat. Plants 1, 1–7 (2015).

71. Zotz, G. et al. Hemiepiphytes revisited. Perspect. Plant Ecol. Evol. Syst. 51, 125620 (2021).

72. Chamberlain, S. et al. rgbif: Interface to the Global Biodiversity Information Facility API. (2022).

73. Nic Lughadha, E., et al. The use and misuse of herbarium specimens in evaluating plant extinction risks. Philos. Trans. R. Soc. B Biol. Sci. 374, 20170402 (2018).

74. Rivers, M. C. et al. How many herbarium specimens are needed to detect threatened species? Biol. Conserv. 144, 2541–2547 (2011).

75. Meyer, C., Weigelt, P. & Kreft, H. Multidimensional biases, gaps and uncertainties in global plant occurrence information. Ecol. Lett. 19, 992–1006 (2016).

76. Zizka, A. et al. No one-size-fits-all solution to clean GBIF. PeerJ 8, e9916 (2020).

77. Brown, M. J. M. et al. rWCVP: a companion R package for the World Checklist of Vascular Plants. New Phytol. **n/a**,.

78. Zizka, A. et al. CoordinateCleaner: Standardized cleaning of occurrence records from biological collection databases. Methods Ecol. Evol. 10, 744–751 (2019).

79. Moat, J. & Bachman, S. rCAT: Conservation Assessment Tools. (2020).

80. Smith, S. A. & Brown, J. W. Constructing a broadly inclusive seed plant phylogeny. Am. J. Bot. 105, 302–314 (2018).

81. Govaerts, R. World Checklist of Vascular Plants (WCVP) – Special Issue 28th February 2022. (2022) doi:10.34885/rar9-jx25.

82. R Core Team. R: A Language and Environment for Statistical Computing. (2021).

83. Stevens, G. C. The Latitudinal Gradient in Geographical Range: How so Many Species Coexist in the Tropics. Am. Nat. 133, 240–256 (1989).

84. Taylor, A., Weigelt, P., König, C., Zotz, G. & Kreft, H. Island disharmony revisited using orchids as a model group. New Phytol. 223, 597–606 (2019).

85. Box, G. E. P. & Cox, D. R. An Analysis of Transformations. J. R. Stat. Soc. Ser. B Methodol. 26, 211–243 (1964).

86. Tung Ho, L. si & Ané, C. A Linear-Time Algorithm for Gaussian and Non-Gaussian Trait Evolution Models. Syst. Biol. 63, 397–408 (2014).

87. Pagel, M. Inferring the historical patterns of biological evolution. Nature 401, 877–884 (1999).

88. Waldron, A. Null Models of Geographic Range Size Evolution Reaffirm Its Heritability. Am. Nat. 170, 221–231 (2007).

89. Villemereuil, P. de, Wells, J. A., Edwards, R. D. & Blomberg, S. P. Bayesian models for comparative analysis integrating phylogenetic uncertainty. BMC Evol. Biol. 12, 102 (2012).

