## Supplementary Material for "A global study of the geographic range size of epiphytes"

### Supporting information

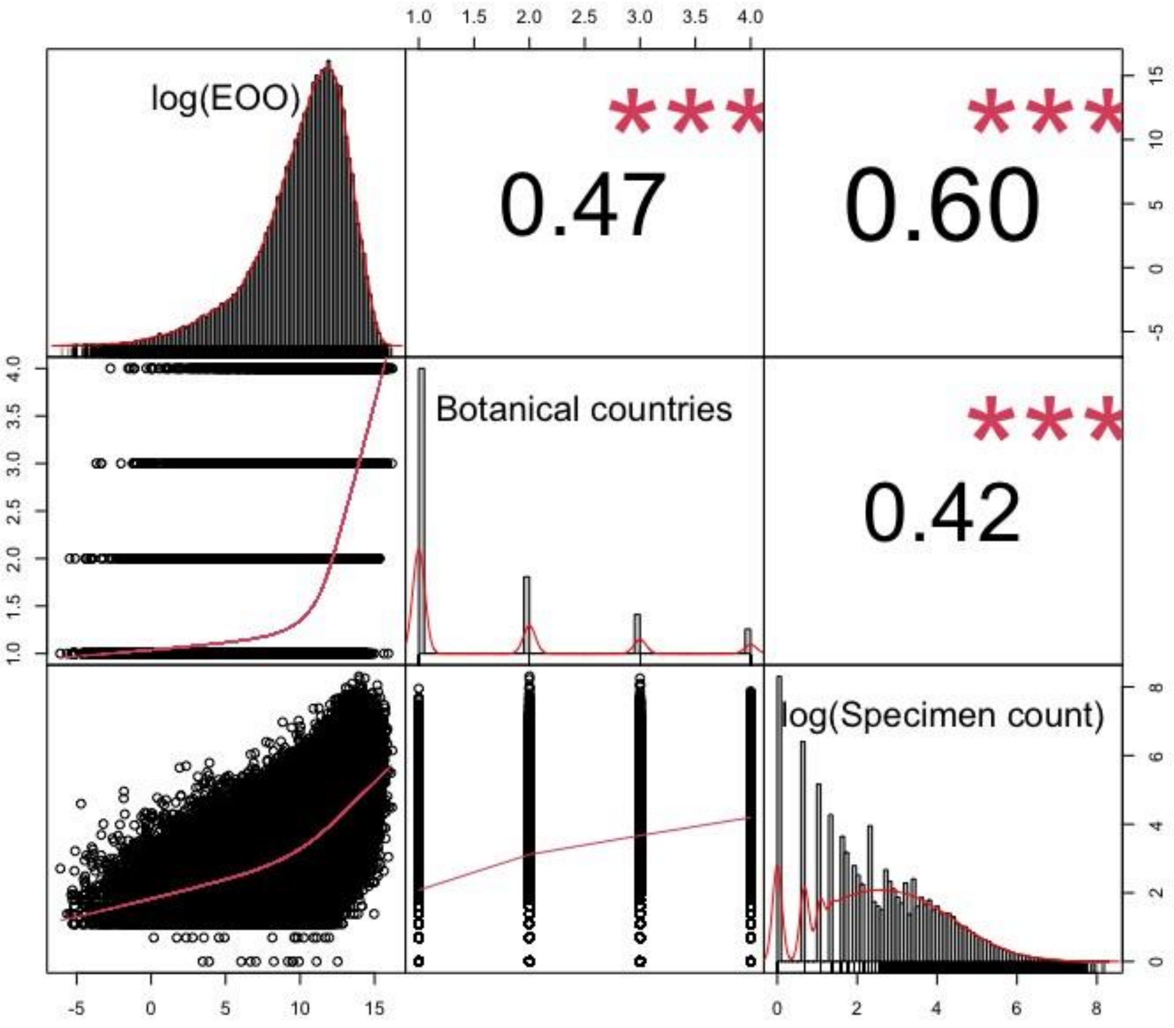


Figure S1: Distribution of individual variables (diagonal), corelation coefficients (top), and corelation plots (bottom) of the three metrics used to measure range size: log(extent of occurrence), botanical countries, and log(specimen count).

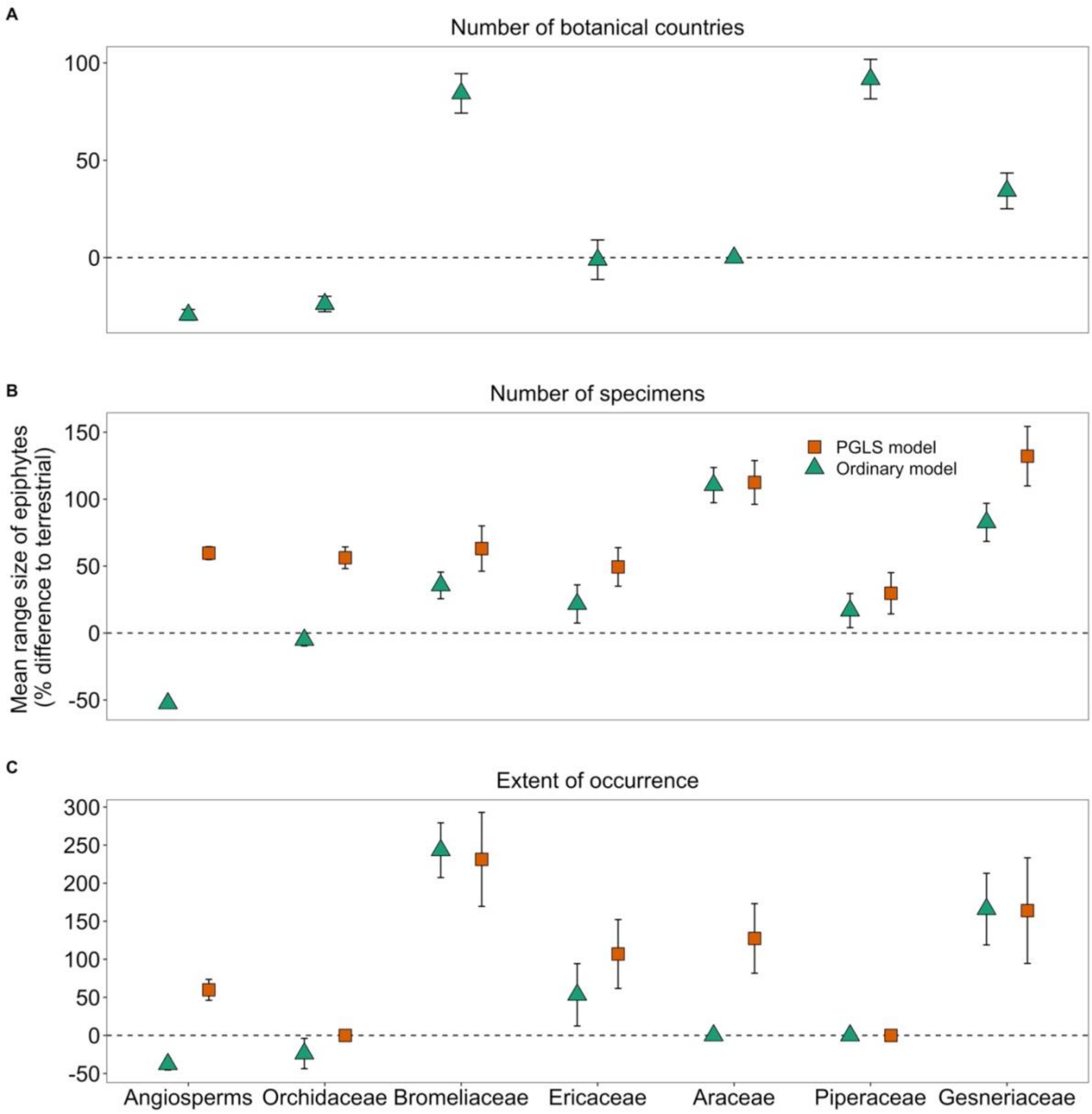


Figure S2: Summary of results for sensitivity analysis restricted to species which have which have the mean latitude of their distribution, measured as botanical countries, in the tropics (between latitudes 23.3 N and 23.3 S). Mean range size is given as percentage (%) difference to terrestrial species with 95% confidence intervals for the three metrics A) number of botanical countries (PGLS not performed); B) specimen count; C) extent of occurrence. The horizontal dotted line represents the baseline of no significant difference in the percentage difference in range size between epiphytes and terrestrial plants.

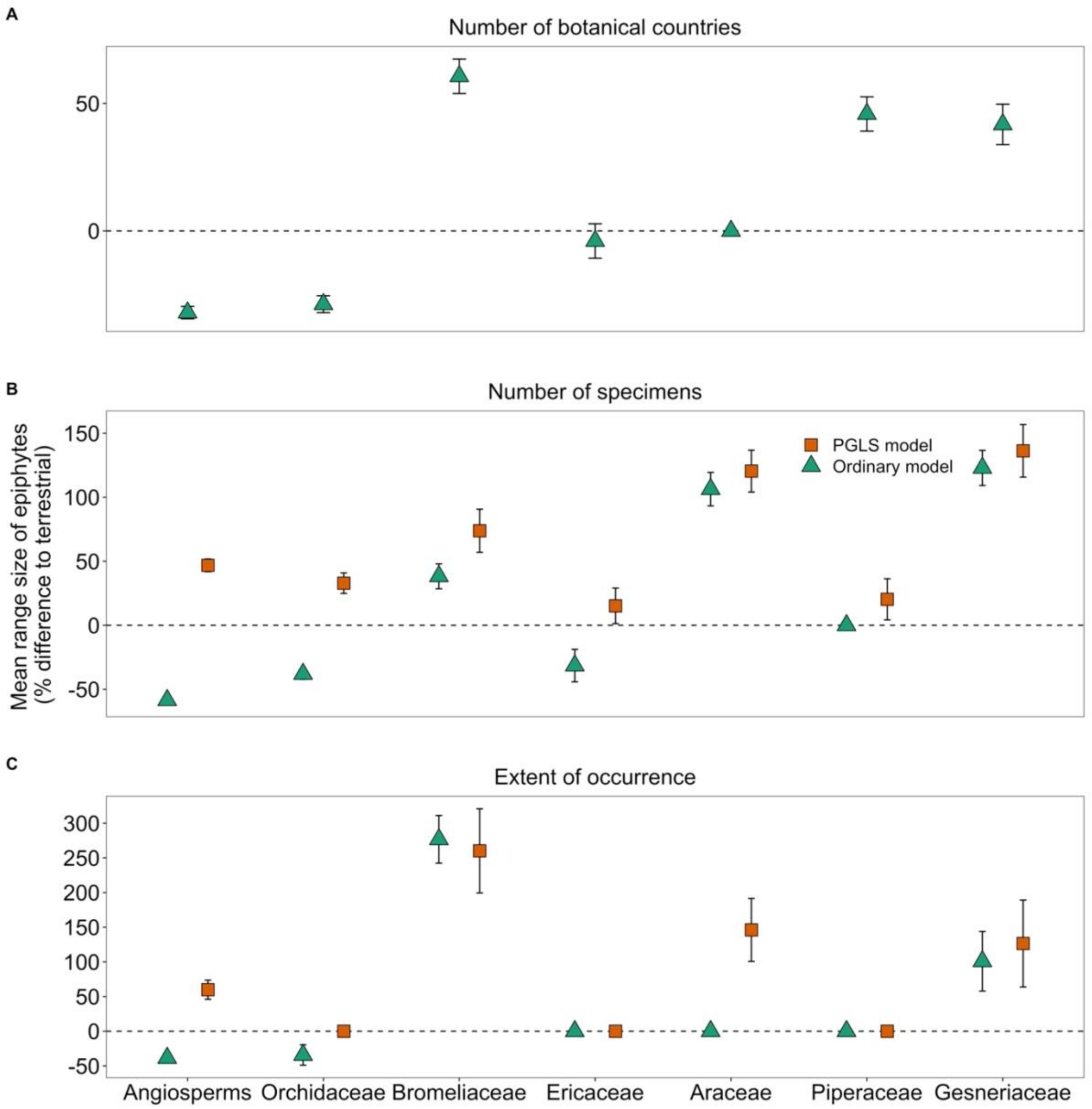


Figure S3: Summary of sensitivity analysis restricted to species which do not occur on any of the 55 botanical countries which are composed of one or more oceanic islands. Mean range size is given as percentage (%) difference to terrestrial species with 95% confidence intervals for the three metrics A) number of botanical countries (PGLS not performed); B) specimen count; C) extent of occurrence. The horizontal dotted line represents the baseline of no significant difference in the percentage difference in range size between epiphytes and terrestrial plants.

Table S1: Estimated effects of epiphytism on range size measured as $\log_{10}$(Specimen count) according to ordinary linear regression models and phylogenetic generalised least squares regression models for angiosperms and each family with more than ten epiphyte species. Coefficients show estimated difference to $\log_{10}$(Specimen count) of terrestrial species.

| | Ordinary regression | | | | Phylogenetic generalised least squares | | | |
|---|---|---|---|---|---|---|---|---|
| Group | Coef. | SE | t-value | p-value | Coef. | SE | t-value | p-value |
| Angiosperms | **-0.867** | **0.011** | **-76.56** | **< 0.001** | **0.38** | **0.025** | **15.216** | **< 0.001** |
| | | | | | **( ± 0.013)** | **( ± < 0.001)** | **( ± 0.497)** | **( ± < 0.001)** |
| Orchidaceae | **-0.456** | **0.020** | **-22.237** | **< 0.001** | **0.286** | **0.041** | **6.966** | **< 0.001** |
| | | | | | **( ± 0.039)** | **( ± 0.001)** | **( ± 0.889)** | **( ± < 0.001)** |
| Bromeliaceae | **0.312** | **0.048** | **6.468** | **< 0.001** | **0.534** | **0.084** | **6.385** | **< 0.001** |
| | | | | | **( ± 0.009)** | **( ± 0.001)** | **( ± 0.135)** | **( ± < 0.001)** |
| Ericaceae | **-0.388** | **0.062** | **-6.218** | **< 0.001** | 0.129 | 0.068 | 1.895 | 0.076 |
| | | | | | ( ± 0.024) | ( ± < 0.001) | ( ± 0.36) | ( ± 0.101) |
| Araceae | **0.723** | **0.064** | **11.242** | **< 0.001** | **0.788** | **0.08** | **9.845** | **< 0.001** |
| | | | | | **( ± 0.007)** | **( ± < 0.001)** | **( ± 0.082)** | **( ± < 0.001)** |
| Piperaceae | **0.183** | **0.062** | **2.930** | **0.003** | **0.199** | **0.065** | **3.044** | **0.003** |
| | | | | | **( ± 0.028)** | **( ± 0.005)** | **( ± 0.18)** | **( ± 0.001)** |
| Gesneriaceae | **0.762** | **0.067** | **11.422** | **< 0.001** | **0.858** | **0.101** | **8.51** | **< 0.001** |
| | | | | | **( ± 0.027)** | **( ± 0.001)** | **( ± 0.256)** | **( ± < 0.001)** |
| Melastomataceae. | 0.140 | 0.08 | 1.748 | 0.081 | **0.59** | **0.097** | **6.061** | **< 0.001** |
| | | | | | **(± 0.024)** | **(± 0.001)** | **(± 0.253)** | **(± < 0.001)** |
| Apocynaceae | **-0.625** | **0.118** | **-5.301** | **< 0.001** | **0.338** | **0.14** | **2.408** | **0.018** |
| | | | | | **(± 0.026)** | **(± 0.001)** | **(± 0.182)** | **(± 0.01)** |
| Rubiaceae | **-0.563** | **0.117** | **-4.833** | **< 0.001** | -0.338 | 0.15 | -2.272 | 0.053 |
| | | | | | (± 0.091) | (± 0.006) | (± 0.663) | (± 0.07) |
| Cactaceae | **0.349** | **0.138** | **2.528** | **0.012** | 0.269 | 0.23 | 1.173 | 0.257 |
| | | | | | (± 0.059) | (± 0.007) | (± 0.273) | (± 0.096) |
| Cyclanthaceae | **0.454** | **0.205** | **2.217** | **0.028** | **0.575** | **0.223** | **2.529** | **0.019** |
| | | | | | **(± 0.178)** | **(± 0.027)** | **(± 0.45)** | **(± 0.012)** |
| Begoniaceae | **1.408** | **0.233** | **6.052** | **< 0.001** | **1.393** | **0.235** | **5.933** | **< 0.001** |
| | | | | | **(± 0.037)** | **(± 0.001)** | **(± 0.161)** | **(± < 0.001)** |

| | | | | | | | | |
|---|---|---|---|---|---|---|---|---|
| Araliaceae | -0.366 | 0.376 | -0.972 | 0.331 | **0.793 (± 0.0244)** | **0.346 (± 0.003)** | **2.290 (± 0.077)** | **0.023 (± 0.005)** |
| Balsaminaceae | 0.068 | 0.228 | 0.300 | 0.764 | 0.072 (± 0.002) | 0.228 (± < 0.001) | 0.318 (± 0.009) | 0.751 (± 0.007) |
| Urticaceae | 0.644 | 0.332 | 1.939 | 0.053 | **0.752 (± 0.028)** | **0.311 (± 0.002)** | **2.418 (± 0.098)** | **0.016 (± 0.005)** |
| Asteraceae | **0.645** | **0.273** | **2.368** | **0.018** | 0.607 (± 0.224) | 0.316 (± 0.021) | 1.919 (± 0.711) | 0.118 (± 0.169) |
| Solanaceae | 0.169 | 0.315 | 0.536 | 0.592 | **0.814 (± 0.081)** | **0.365 (± 0.007)** | **2.226 (± 0.197)** | **0.029 (± 0.02)** |
| Crassulaceae | **0.868** | **0.301** | **2.883** | **0.004** | **1.00 (± 0.025)** | **0.292 (± 0.001)** | **3.418 (± 0.09)** | **0.001 (± < 0.001)** |
| Campanulaceae | **0.987** | **0.278** | **3.544** | **< 0.001** | **1.458 (± 0.064)** | **0.301 (± 0.004)** | **4.839 (± 0.213)** | **< 0.001 (± < 0.001)** |
| Zingiberaceae | 0.4 | 0.269 | 1.485 | 0.138 | **0.69 (± 0.028)** | **0.274 (± 0.002)** | **2.52 (± 0.108)** | **0.012 (± 0.004)** |
| Nepenthaceae | 0.488 | 0.345 | 1.416 | 0.16 | 0.492 (± 0.101) | 0.342 (± 0.007) | 1.446 (± 0.324) | 0.172 (± 0.09) |
| Lentibularaceae. | **0.897** | **0.385** | **2.327** | **0.021** | **0.881 (± 0.022)** | **0.385 (± 0.001)** | **2.286 (± 0.058)** | **0.023 (± 0.004)** |
| Asparagaceae | 0.457 | 0.421 | 1.086 | 0.278 | 0.218 (± 0.066) | 0.382 (± 0.003) | 0.571 (± 0.171) | 0.573 (± 0.119) |
| Schlegeliaceae | -0.239 | 0.465 | -0.513 | 0.612 | -0.239 (± < 0.001) | 0.465 (± < 0.001) | -0.513 (± < 0.001) | 0.612 (± < 0.001) |

Table S2: Estimated effects of epiphytism on range size measured as $\log_{10}$(Extent of occurrence) according to ordinary linear regression models and phylogenetic generalised least squares regression models for angiosperms and each family with more than ten epiphyte species. Coefficients show estimated difference to $\log_{10}$(Extent of occurrence) of terrestrial species.

| | Ordinary regression | | | | Phylogenetic generalised least squares | | | |
|---|---|---|---|---|---|---|---|---|
| Group | Coef. | SE | t-value | p-value | Coef. | SE | t-value | p-value |
| Angiosperms | **-0.402** | **0.035** | **-11.509** | **<0.001** | **0.434** | **0.068** | **6.379** | **<0.001** |
| | | | | | **(± 0.02)** | **(± <0.001)** | **(± 0.288)** | **(± <0.001)** |
| Orchidaceae | **-0.39** | **0.07** | **-5.547** | **<0.001** | 0.087 | 0.14 | 0.627 | 0.525 |
| | | | | | (± 0.073) | (± 0.003) | (± 0.525) | (± 0.25) |
| Bromeliaceae | **1.32** | **0.161** | **8.221** | **<0.001** | **1.284** | **0.269** | **4.765** | **<0.001** |
| | | | | | **(± 0.037)** | **(± 0.002)** | **(± 0.166)** | **(± <0.001)** |
| Ericaceae | 0.201 | 0.146 | 1.372 | 0.17 | 0.017 | 0.171 | 0.099 | 0.884 |
| | | | | | (± 0.028) | (± 0.001) | (± 0.161) | (± 0.092) |
| Araceae | 0.079 | 0.163 | 0.485 | 0.628 | **0.873** | **0.208** | **4.19** | **<0.001** |
| | | | | | **(± 0.018)** | **(± 0.001)** | **(± 0.084)** | **(± <0.001)** |
| Piperaceae | -0.271 | 0.192 | -1.412 | 0.158 | -0.227 | 0.2 | -1.167 | 0.289 |
| | | | | | (± 0.082) | (± 0.014) | (± 0.452) | (± 0.242) |
| Gesneriaceae | **0.940** | **0.200** | **4.708** | **<0.001** | **0.883** | **0.278** | **3.173** | **0.002** |
| | | | | | **(± 0.044)** | **(± 0.004)** | **(± 0.125)** | **(± 0.001)** |
| Melastomataceae. | **0.513** | **0.224** | **2.289** | **0.022** | **1.342** | **0.318** | **4.221** | **<0.001** |
| | | | | | **(± 0.062)** | **(± 0.003)** | **(± 0.198)** | **(± <0.001)** |
| Apocynaceae | 0.008 | 0.547 | 0.014 | 0.989 | 0.512 | 0.633 | 0.809 | 0.425 |
| | | | | | (± 0.106) | (± 0.006) | (± 0.165) | (± 0.099) |
| Rubiaceae | 0.017 | 0.363 | 0.046 | 0.963 | 0.184 | 0.392 | 0.469 | 0.652 |
| | | | | | (± 0.126) | (± 0.004) | (± 0.321) | (± 0.191) |
| Cactaceae | 0.18 | 0.35 | 0.515 | 0.606 | 0.242 | 0.552 | 0.439 | 0.666 |
| | | | | | (± 0.096) | (± 0.012) | (± 0.178) | (± 0.126) |
| Cyclanthaceae | -0.461 | 0.425 | -1.084 | 0.280 | -0.458 | 0.426 | -1.076 | 0.285 |
| | | | | | (± 0.028) | (± 0.011) | (± 0.075) | (± 0.046) |
| Begoniaceae | 1.109 | 0.607 | 1.829 | 0.068 | 1.11 | 0.608 | 1.825 | 0.069 |

| | | | | | | | | |
|---|---|---|---|---|---|---|---|---|
| | | | | | (± 0.027) | (± 0.002) | (± 0.045) | (± 0.007) |
| Araliaceae | 0.311 | 0.948 | 0.328 | 0.743 | 0.784 | 0.980 | 0.801 | 0.425 |
| | | | | | (± 0.073) | (±0.005) | (±0.074) | (±0.042 |
| Balsaminaceae | -0.125 | 0.944 | -0.132 | 0.895 | -0.098 | 0.945 | -0.103 | 0.909 |
| | | | | | (± 0.057) | (± 0.004) | (± 0.061) | (± 0.026) |
| Urticaceae | 1.634 | 1.079 | 1.515 | 0.130 | 1.977 | 1.059 | 1.866 | 0.063 |
| | | | | | (± 0.054) | (± 0.002) | (± 0.051) | (± 0.007) |
| Asteraceae | 0.560 | 0.588 | 0.952 | 0.341 | 0.764 | 0.679 | 1.121 | 0.309 |
| | | | | | (± 0.336) | (± 0.041) | (± 0.485) | (± 0.223) |
| Solanaceae | 0.104 | 0.606 | 0.171 | 0.864 | 0.064 | 0.713 | 0.091 | 0.918 |
| | | | | | (± 0.063) | (± 0.015) | (± 0.091) | (± 0.06) |
| Crassulaceae | -0.193 | 0.634 | -0.304 | 0.761 | 0.24 | 0.612 | 0.393 | 0.696 |
| | | | | | (± 0.058) | (± 0.003) | (± 0.095) | (± 0.07) |
| Campanulaceae. | -0.641 | 0.610 | -1.051 | 0.293 | 1.23 | 0.659 | 1.863 | 0.086 |
| | | | | | (± 0.264) | (± 0.014) | (± 0.393) | (± 0.098) |
| Zingiberaceae | 0.200 | 0.867 | 0.231 | 0.818 | 0.2 | 0.867 | 0.231 | 0.818 |
| | | | | | (± <0.001) | (± <0.001) | (± <0.001) | (± <0.001) |
| Nepenthaceae | 0.016 | 0.997 | 0.016 | 0.987 | 0.016 | 0.997 | 0.016 | 0.987 |
| | | | | | (± <0.001) | (± <0.001) | (± <0.001) | (± <0.001) |
| Lentibularaceae. | **2.164** | **1.081** | **2.003** | **0.047** | 1.96 | 1.077 | 1.82 | 0.073 |
| | | | | | (± 0.135) | (± 0.01) | (± 0.128) | (± 0.018) |
| Asparagaceae | 0.727 | 0.951 | 0.764 | 0.445 | 0.065 | 0.922 | 0.07 | 0.934 |
| | | | | | (± 0.094) | (± 0.002) | (± 0.102) | (± 0.071) |
| Schlegeliaceae | **-3.397** | **0.879** | **-3.864** | **0.001** | **-3.374** | **0.889** | **-3.811** | **0.002** |
| | | | | | **(± 0.119)** | **(± 0.048)** | **(± 0.265)** | **(± 0.004)** |

Table S3: Estimated effects of epiphytism on range size measured as Number of botanical countries according to generalised linear models with quasipoisson error distributions for angiosperms and each family with more than ten epiphyte species. Coefficients show estimated difference to number of botanical countries of terrestrial species.

| | Generalised linear regression | | | |
|---|---|---|---|---|
| Group | Coef. | SE | t-value | p-value |
| Angiosperms | **-0.443** | **0.015** | **-29.98** | **<0.001** |
| Orchidaceae | **-0.377** | **0.018** | **-20.4** | **<0.001** |
| Bromeliaceae | **0.595** | **0.047** | **12.548** | **<0.001** |
| Ericaceae | **-0.302** | **0.111** | **-2.727** | **0.006** |
| Araceae | -0.207 | 0.118 | -1.746 | 0.081 |
| Piperaceae | **0.634** | **0.056** | **11.221** | **<0.001** |
| Gesneriaceae | **0.375** | **0.039** | **9.554** | **<0.001** |
| Melastomataceae | -0.135 | 0.079 | -1.713 | 0.087 |
| Apocynaceae | -0.057 | 0.100 | -0.571 | 0.568 |
| Rubiaceae | -0.027 | 0.129 | -0.214 | 0.831 |
| Cactaceae | **0.657** | **0.0800** | **8.251** | **<0.001** |
| Cyclanthaceae | 0.171 | 0.154 | 1.109 | 0.269 |
| Begoniaceae | **1.142** | **0.08** | **14.208** | **<0.001** |
| Araliaceae | 0.262 | 0.246 | 1.068 | 0.286 |
| Balsaminaceae | -0.082 | 0.307 | -0.266 | 0.79 |
| Urticaceae | **0.938** | **0.209** | **4.485** | **<0.001** |
| Asteraceae | 0.246 | 0.266 | 0.924 | 0.356 |
| Solanaceae | 0.077 | 0.302 | 0.255 | 0.799 |
| Crassulaceae | 0.115 | 0.287 | 0.401 | 0.688 |
| Campanulaceae. | -0.388 | 0.396 | -0.98 | 0.327 |
| Zingiberaceae | 0.345 | 0.189 | 1.822 | 0.069 |
| Nepenthaceae | -0.234 | 0.236 | -0.989 | 0.324 |
| Lentibularaceae. | 0.093 | 0.411 | 0.227 | 0.820 |
| Asparagaceae | 0.065 | 0.333 | 0.196 | 0.845 |

| | | | | |
|---|---|---|---|---|
| Schlegeliaceae | -0.157 | 0.323 | -0.485 | 0.631 |

Table S4: Summary of the number and proportion of epiphytic species in angiosperms and each of the twenty-four angiosperm families with 10 or more epiphytic species.

| Group | Number of epiphytes | Epiphyte proportion |
|---|---|---|
| Angiosperms | 27,544 | 0.81 |
| Orchidaceae | 21,034 | 0.694 |
| Bromeliaceae | 1,937 | 0.546 |
| Ericaceae | 876 | 0.192 |
| Araceae | 766 | 0.186 |
| Piperaceae | 699 | 0.183 |
| Gesneriaceae | 644 | 0.171 |
| Melastomataceae . | 362 | 0.063 |
| Apocynaceae | 247 | 0.038 |
| Rubiaceae | 189 | 0.013 |
| Cactaceae | 138 | 0.077 |
| Cyclanthaceae | 60 | 0.26 |
| Begoniaceae | 59 | 0.031 |
| Araliaceae | 54 | 0.033 |
| Balsaminaceae | 47 | 0.044 |
| Urticaceae | 41 | 0.041 |
| Asteraceae | 39 | 0.001 |
| Solanaceae | 35 | 0.013 |
| Crassulaceae | 34 | 0.020 |
| Campanulaceae | 33 | 0.013 |
| Zingiberaceae | 33 | 0.018 |
| Nepenthaceae | 30 | 0.163 |

| | | |
|---|---|---|
| Lentibularaceae | 28 | 0.067 |
| Asparagaceae | 25 | 0.008 |
| Schlegeliaceae | 16 | 0.432 |

Table S5: Summary of GBIF download citations.

| |
|---|
| GBIF.org (21 March 2022) GBIF Occurrence Download https://doi.org/10.15468/dl.wmerwv |
| GBIF.org (21 March 2022) GBIF Occurrence Download https://doi.org/10.15468/dl.uj3k3y |
| GBIF.org (21 March 2022) GBIF Occurrence Download https://doi.org/10.15468/dl.95tcrx |
| GBIF.org (21 March 2022) GBIF Occurrence Download https://doi.org/10.15468/dl.t22czn |
| GBIF.org (23 March 2022) GBIF Occurrence Download https://doi.org/10.15468/dl.nje5yj |
| GBIF.org (27 March 2022) GBIF Occurrence Download https://doi.org/10.15468/dl.u9zmqc |

Table S6: Number and percentage of species which occur in four or fewer botanical countries included in each of the two GBIF-derived datasets (EOO and specimen count). We chose to tabulate percentage of species per group which occur in four or fewer botanical countries rather than percentage of total number of species per group in order to better illustrate data coverage and potential biases between groups in GBIF data, given that we did not attempt to download GBIF data for the 16% of angiosperm species found in five or more botanical countries.

| | Number of species in EOO analysis | | | Number of species in Specimen count analysis | | |
|---|---|---|---|---|---|---|
| | All species | Epiphytes | Terrestrials | All species | Epiphytes | Terrestrials |
| Angiosperms | 145,898 (51%) | 8,170 (33%) | 137,728 (53%) | 244,461 (86%) | 20,326 (82%) | 224,135 (87%) |
| Orchidaceae | 7,634 (28%) | 4,811 (25%) | 2,823 (36%) | 20,906 (78%) | 14,903 (78%) | 6,003 (76%) |
| Bromeliaceae | 1,719 (52%) | 929 (53%) | 790 (50%) | 30,78 (92%) | 1,614 (93%) | 1,464 (92%) |
| Ericaceae | 2,812 (66%) | 531 (64%) | 2,281 (67%) | 3,833 (90%) | 792 (95%) | 3041 (89%) |
| Araceae | 1,625 (44%) | 506 (72%) | 1,119 (37%) | 3088 (83%) | 685 (97%) | 2,403 (80%) |
| Piperaceae | 1,563 (44%) | 289 (49%) | 1,274 (43%) | 3,214 (90%) | 574 (97%) | 2,640 (89%) |
| Gesneriaceae | 1,490 (41%) | 346 (59%) | 1,144 (38%) | 3,146 (87%) | 539 (92%) | 2,607 (86%) |
